# Effects of host species identity and diet on biodiversity of oral and rectal microbiomes of Puerto Rican bats

**DOI:** 10.1101/2020.01.31.928994

**Authors:** Steven J. Presley, Joerg Graf, Ahmad F. Hassan, Anna R. Sjodin, Michael R. Willig

**Affiliations:** Institute of the Environment, Center for Environmental Sciences & Engineering, and Department of Ecology & Evolutionary Biology, University of Connecticut, Storrs, Connecticut 06269-4210; Department of Molecular & Cell Biology, University of Connecticut, Storrs, Connecticut 06269-3125; Department of Biological Sciences, University of Idaho, Moscow, Idaho 83844

**Keywords:** Chiroptera, Mormoopidae, Phyllostomidae, spatial variation, Vespertilionidae

## Abstract

Microbiomes perform vital functions for their mammalian hosts, making them potential drivers of host evolution. Understanding effects of environmental factors and host characteristics on the composition and biodiversity of microbiomes may provide novel insights into the origin and maintenance of these symbiotic relationships. Our goals were to (1) characterize biodiversity of oral and rectal microbiomes of bats in Puerto Rico; and (2) determine the effects of geographic location and host characteristics on that biodiversity. We collected bats and their microbiomes from 3 sites, and used 4 metrics (species richness, Shannon diversity, Camargo evenness, Berger-Parker dominance) to characterize biodiversity. We evaluated the relative importance of site, host sex, host species identity, and host foraging guild on microbiome biodiversity. Microbiome biodiversity was highly variable among conspecifics. Geographical location exhibited consistent effects, whereas host sex did not do so. Within each host guild, host species exhibited consistent differences in oral and rectal microbiome biodiversity. Oral microbiome biodiversity was indistinguishable between guilds, whereas rectal microbiome biodiversity was significantly greater in carnivores than in herbivores. The high intraspecific and spatial variation in microbiome biodiversity necessitate a large number of samples to isolate the effects of environmental or host characteristics on microbiomes. Species-specific biodiversity of oral microbiomes suggests these communities are structured by direct interactions with the host immune system via epithelial receptors. In contrast, the number of microbial taxa that a host gut supports may be contingent on the number and kinds of functions a host requires of its microbiome.

## INTRODUCTION

Microbiomes perform vital functions for their mammalian hosts, including nutrient acquisition, pathogen defense, and immune development [1–3]. This suggests that microbiomes may be important drivers of host evolution, affecting their physiology, immunocompetence, diet, and ultimately fitness [4]. Moreover, aspects of mammalian physiology, anatomy, behavior, diet, and niche affect which microbes encounter particular host habitats (e.g. skin, oral cavity, gastrointestinal tract). Consequently, these symbiotic associations likely represent co-evolutionary relationships [5, 6].

Understanding effects of environmental factors and host characteristics on the composition and biodiversity of microbiomes may provide novel insights into the origin and maintenance of symbiotic relationships. Indeed, host phylogeny, host diet, and environmental characteristics are the primary candidates likely to affect variation in microbiome composition or biodiversity [7–11]. Host phylogeny is a particularly attractive explanation as it forms the basis for coevolutionary dynamics. Because organisms evolve via descent with modification, phylogenetic inertia gives rise to a priori expectations that more closely related species will be more similar (from genetic, functional, behavioral, anatomical, and physiological perspectives), and that more distantly related species will be less similar [12]. Consequently, host phylogeny may be an effective proxy for combinations of host characteristics that affect the composition and biodiversity of microbiomes rather than an explanatory mechanism for variation among hosts in aspects of their microbiome per se.

Host diet has been a focal point for understanding microbiome composition and biodiversity, especially for gastrointestinal microbiomes, which facilitate digestive processes and are directly exposed to the ingested food. Consequently, intra- or inter-specific differences in host diet may result in differences in gastrointestinal microbiomes due to exposure (i.e. animals with similar diets may consume similar microbiota) or due to the digestive functions provided by the microbiota [9, 13]. In addition, hosts that live in similar environments may be exposed to similar microbiota, resulting in similar microbiome composition or biodiversity [13]. Important aspects of the environment that may affect microbiome composition and biodiversity include host abundance and community composition, as well as ambient environmental characteristic s (e.g. roost type, habitat type, and abiotic factors).

Studies typically consider samples from intestinal contents, intestinal linings, or feces to represent the same microbial communities [9, 14, 15]. Nonetheless, microbiomes isolated from the mucosal layer of the intestines are distinct from those isolated from feces or intestinal contents [14]. Moreover, variation among microbiomes from the intestinal mucosa are closely associated with host evolutionary relationships, whereas variation in fecal microbiomes are closely associated with dietary variation among hosts [14].

### Bats as microbiome hosts

Bats are an ideal model taxon for the study of variation in microbiome biodiversity [14]. They compose the 2^nd^ most species-rich order of mammals, and are nearly cosmopolitan, occurring everywhere but the Arctic, Antarctica, and a few small oceanic islands [16]. In addition, bats are locally abundant, live in close proximity to humans, travel long distances to forage or between winter and summer ranges, and are functionally diverse [17, 18]. Moreover, bats are important agents of pollination, seed dispersal, and pest control [19, 20], and exhibit specializations to forage on nectar, fruit, insects, fish, small vertebrates, and blood [17, 18, 21, 22]. As with other mammals, functional traits and behaviors are evolutionarily conserved in bats, often confounding the ability to evaluate independent effects of diet or phylogeny on ecological patterns [23].

Understanding bat microbiome composition and diversity may be especially important because bats often live in proximity to humans [24] and are reservoirs or vectors for many well-known zoonoses [25–29]. Their presence in settlements affects infection rates of diseases in humans [30]. Bats use many human-dominated habitats: they feed on fruits in orchards, forage for insects around streetlights, and use buildings for maternity colonies, roosts, and hibernacula [24]. In addition, bats are highly vagile and capable of traveling long distances in a single night. This creates opportunities for exposure to novel microbes and to enhance dispersal [31]. In addition, microbiomes may drive host evolution, physiology, and fitness [2]. For example, the successful evolution of new dietary strategies within a clade (e.g. the diversification of herbivorous strategies within the Phyllostomidae) may be contingent of the functional diversity of their associated microbiomes. Finally, the digestive physiology of flying vertebrates (bats and birds) differ from that of other vertebrates [32], including reliance on paracellular glucose absorption, resulting in different mechanisms structuring the microbiomes of bats and birds than in terrestrial vertebrate groups.

Thirteen species of bat occur on Puerto Rico [33], including 7 insectivores (*Eptesicus fuscus*, *Lasiurus borealis*, *Molossus molossus*, *Mormoops blainvillii*, *Pteronotus quadridens*, *P*. *parnellii*, *Tadarida brasiliensis*), a piscivore (*Noctilio leporinus*), a nectarivore (*Monophyllus redmani*), 2 frugivores (*Artibeus jamaicensis*, *Stenoderma rufum*), and 2 generalist herbivores (*Brachyphylla cavernarum*, *Erophylla sezekorni*). Bats that consume fruit, nectar, flowers, or pollen typically have diverse herbivorous diets that differ in preferred dietary items, with *M*. *redmani* being primarily nectarivorous, *A*. *jamaicensis* and *S*. *rufum* being primarily frugivorous, and *E*. *sezekorni* and *B*. *cavernarum* being generalist herbivores [33]. The insectivores belong to 3 families (Vespertilionidae, Molossidae, and Mormoopidae); the piscivore is a noctilionid; and phyllostomids consume fruit, flowers, pollen, or nectar. The Noctilionidae, Mormoopidae, and Phyllostomidae are members of the Noctilionoidea superfamily, whereas the Vespertilionidae and Molossidae are members of the Vespertilionoidea superfamily [34]. These systematic relationships decouple insectivory from phylogeny and may help disentangle the relative effects of evolutionary history and ecological function as drivers of microbiome composition and diversity. We grouped bats into broad foraging guilds (carnivores and herbivores) to evaluate effects of general diet on microbiome biodiversity.

### Host environments

Oral microbiomes provide benefits to the host, including prevention of infection by exogenous microorganisms, regulation of immune responses, and the conversion of dietary nitrates into nitrites that improve vascular health and stimulate gastric mucus production [35, 36]. The oral environment (e.g. pH, saliva, temperature, nutrient sources, aerobic conditions) determines which microbes colonize and become minor or major components of the oral microbiome [37]. In addition, the microbiome can modify the environment, facilitating or preventing establishment by other microbes. Despite the importance of oral microbiomes to their hosts, they have rarely been studied in wild animals (but see [15]) even though such studies are critical for advancing evolutionary ecology in general [38].

Studies rarely have sufficient sample sizes from multiple locations, species, or foraging guilds to powerfully and simultaneously address multiple factors that affect microbiome composition or biodiversity in bats (but see [15]). Moreover, studies are lacking that simultaneous consider effects of environmental and host factors on microbiomes from multiple sources (e.g. oral cavity). To address these issues, we collected oral and rectal samples from bats captured at three locations (hereafter called “sites”) in Puerto Rico. We evaluated the relative importance of site, host sex, host species identity, and host foraging guild on microbiome biodiversity from oral and rectal samples. We used a hierarchical analytical design to evaluate these factors (Fig. 1). First, for each host species with sufficient sample sizes from multiple caves, we evaluated effects of site (i.e. host population) and host sex on microbiome biodiversity. Second, we evaluated the effect of host species identity on microbiome biodiversity separately for bats within each of two broadly defined foraging guilds. Finally, we evaluated the effect of host foraging guild (carnivores versus herbivores) on microbiome biodiversity.

**Fig. 1.**
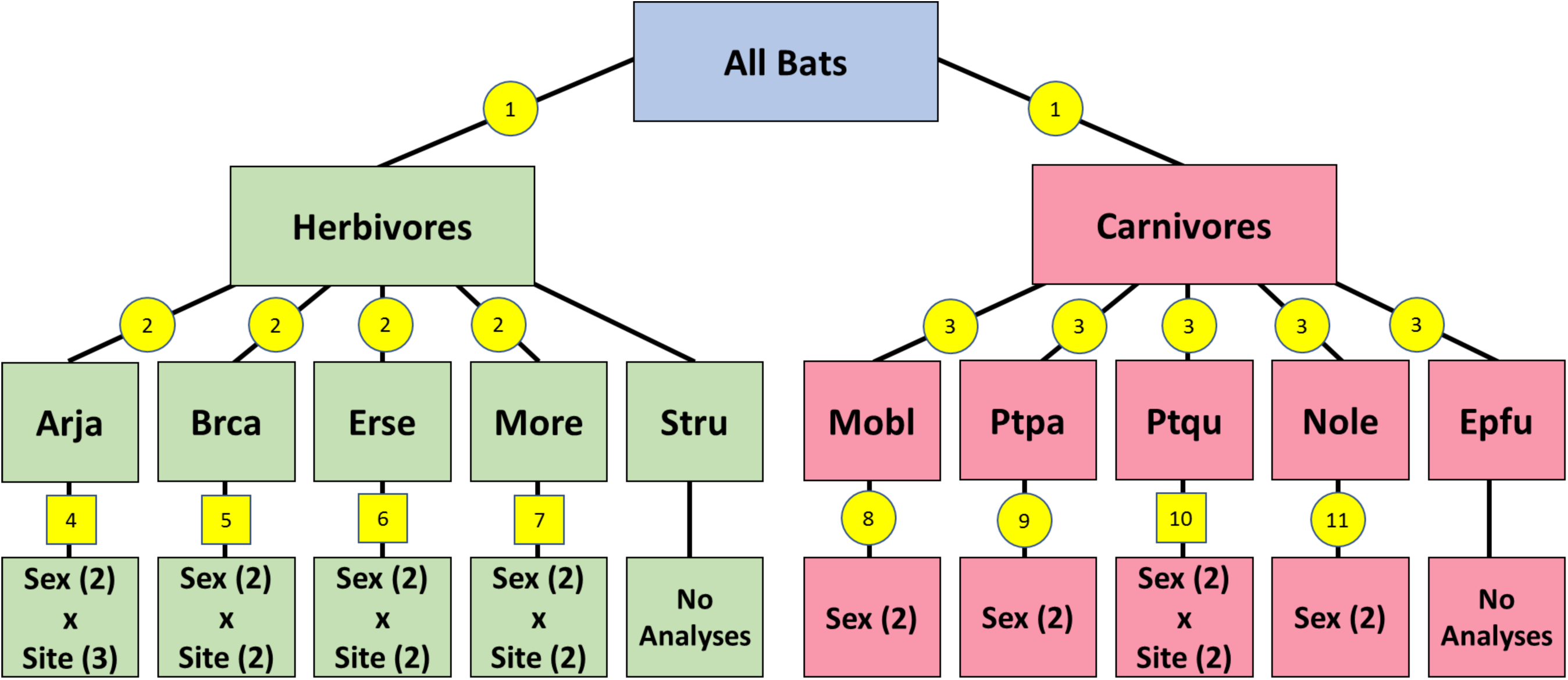
Hierarchical design of statistical analyses. Yellow shapes indicate analyses: circle, General Linear Mixed-Effects Model (GLMM); square, Analysis of Variance (ANOVA). Numbers indicate particular statistical analyses that compare groups: 1, guilds; 2 herbivorous species; 3, carnivorous species; and 4-11, combinations of sex and cave or only sex for each species. Numbers in parentheses equal the number of treatment levels in a factor. Abbreviations are: Arja, *Artibeus jamaicensis*; Brca, *Brachyphylla cavernarum*; Erses, *Erophylla sezekorni*; More, *Monophyllus redmani*; Stru, *Stenoderma rufum*; Mobl, *Mormoops blainvillii*; Ptpa, *Pteronotus parnellii*; Ptqu, *Pteronotus quadridens*; Nole, *Noctilio leporinus*; and Epfu, *Eptesicus fuscus*. Only 1 sample was collected from *S.rufum*; therefore, this species was omitted from the interspecific comparison within the herbivore guild.

We expected factors that mold patterns in oral microbiomes to be different from those that mold such patterns in rectal microbiomes. More specifically, we expected dietary guild to have a larger impact on the biodiversity of rectal microbiomes than on that of oral microbiomes because sources of nutrients and energy (fats, carbohydrates, proteins) have a dominant effect on the composition and diversity of microbiomes associated with the digestive tract [8]. In contrast, we expected biodiversity of the oral microbiome, but not that of the rectal microbiome, to respond to host species identity and geographical site because oral microbiomes are affected primarily by the interactions with the epithelia and exposure to local habitats (e.g. roost locations, animals that share a roost, hot cave versus cold cave).

## METHODS

### Study area and sample collection

Field work was conducted at three sites (Mata de Plátano, Río Encantado, and Aguas Buenas) in Puerto Rico (Fig. 2), Each is in an area characterized by limestone formations (karst region), in which weathering has produced ridges, towers, fissures, sinkholes, and caves throughout the landscape. Although bats captured in a location may not be roosting in a single cave, all are using the same habitats and resources, meeting the criteria for a population.

**Fig. 2.**
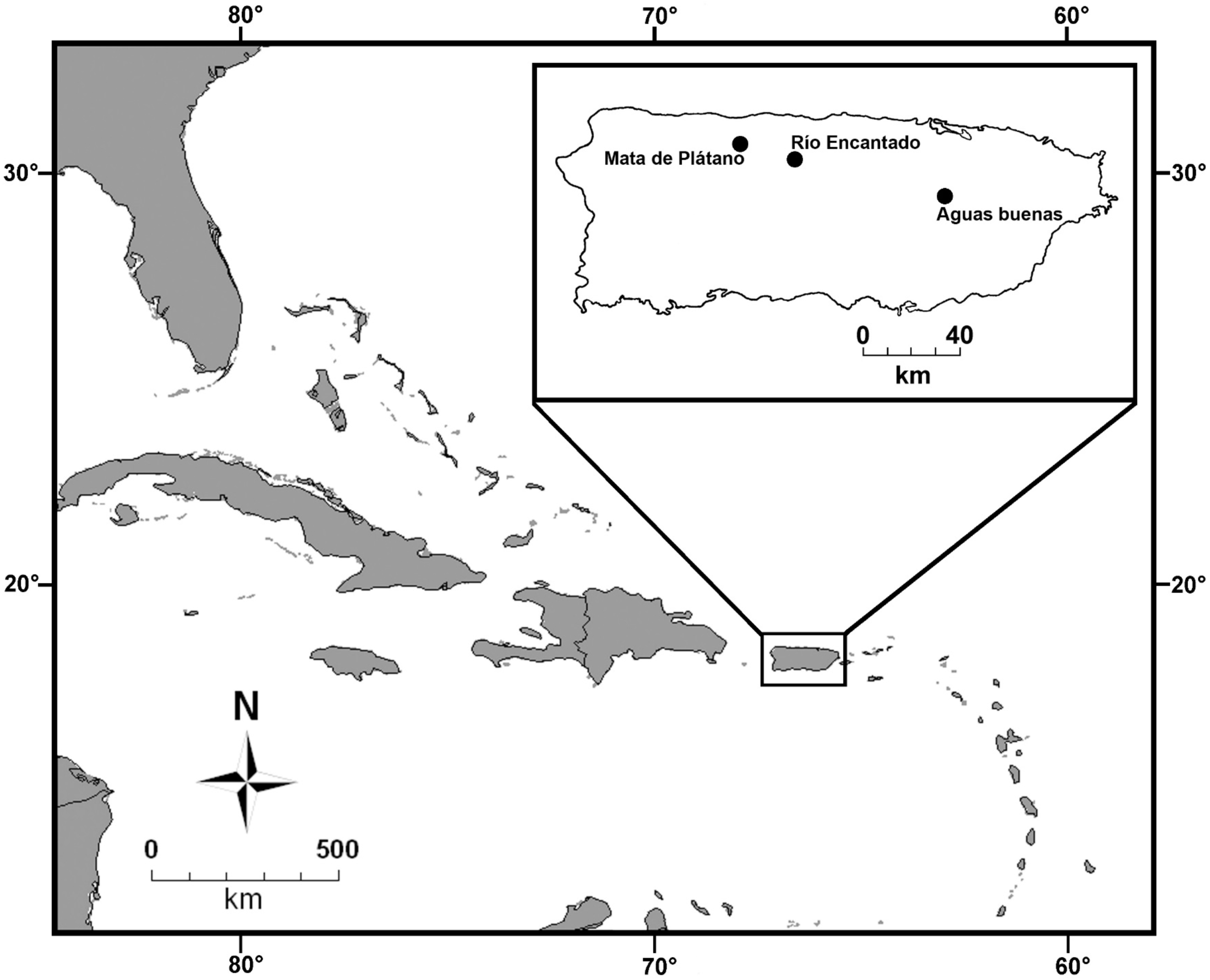
Map of the Caribbean showing the location of Puerto Rico within the Antilles as well as the location of the three collection localities in Puerto Rico.

The majority of sampling was conducted on the Mata de Plátano Nature Reserve (operated by InterAmerican University, Bayamon, Puerto Rico) in north-central Puerto Rico (18° 24.87’ N, 66° 43.53’ W). Mata de Plátano harbors two adjacent, well-studied caves (Culebrones and Larva). Culebrones is a structurally complex hot cave, with temperatures reaching 40 °C and relative humidity at 100%. It is home to about 300,000 bats representing six species [39]: three carnivores (*P quadridens*, *P*. *parnellii*, *M*. *blainvillii*) and three herbivores (*M*. *redmani*, *E*. *sezekorni*, *B*. *cavernarum*). Bats were sampled at Culebrones for 28 nights from June to August 2017. A harp trap was placed at sunset immediately outside the cave opening and monitored continually until the maximum number of bats that could be processed in a single night was captured. The harp trap was used at Culebrones because the cave has a single, small opening, that funnels hundreds of thousands of bats through a small space as bats emerge during and after sunset.

Larva is a cold cave that is much smaller, cooler, and less structurally complex than Culebrones. Only a small number of bats (30-200) representing two species (*A*. *jamaicensis* and *E*. *fuscus*) roost in the cave. Bats were sampled from Larva on seven occasions from June to August of 2017, using two different techniques. After sunset, mist nets were placed along a trail outside of the cave entrance and were checked at least every ten minutes. Because few individuals were captured with mist nets, hand nets were used to capture bats inside the cave to increase sample sizes.

Río Encantado is home to Ramon Cave (18° 21.41’ N, 66° 32.36’ W), a large, cool cave known to support a single bat species, *A. jamaicensis* [33]. The cave is 10 km southeast of Mata de Plátano and is associated with an extensive underground river system. The underground river has many openings throughout its range, but only a single opening exists at this location. Habitats surrounding Ramon Cave are owned and protected by a non-profit organization (Para la Naturaleza). Bats were sampled at Río Encantado on six nights during July of 2017. A harp trap was placed near the cave entrance and mist nets were placed along the trail leading to the cave. Harp traps were monitored continually, and mist nets were checked at least every 10 minutes. Bats were captured from sunset until the maximum number of bats that could be processed in a single night was collected.

Aguas Buenas is a cool cave that is located 70 km southeast of Mata de Plátano (18° 14.01’ N, 66° 6.30’ W). *Artibeus jamaicensis*, *B. cavernarum*, *M. redmani*, *P. quadridens*, *E. fuscus*, and *L*. *borealis* have been recorded roosting in or flying near the cave [33]. Bats were captured at Aguas Buenas on four nights in July and August of 2017. The entrance to the cave is not easily accessible, as it is elevated above ground-level and blocked by a river. Consequently, bats were captured using mist nets at each of the two major flyways from the cave: along the trail to the cave and across the river outside of the cave. Nets were opened at sunset and monitored at least every 10 minutes until approximately 01:00 or until the maximum number of bats that could be processed in a single night was collected.

Species identity, sex, reproductive status and mass were determined for each captured individual prior to placement in a cotton holding bag. Separate, clean cotton-tipped swabs were used to collect saliva from the mouth or feces from the rectum and anal region of each bat. Swabs were placed in individual cryovials and sent to the University of Connecticut at −80 °C in a dry ice shipper. All methods were approved by the University of Connecticut Institutional Animal Care and Use Committee (IACUC, protocol A15-032).

### Microbiome Analysis

DNA was extracted using DNeasy PowerSoil kit (Qiagen). Swabs were shaved off to maximize DNA output using sterile surgical blades, carbon steel, Size 15 (Bard-Parker). The DNeasy PowerSoil protocol was followed, but instead of vortexing the bead tubes a PowerLyzer 24 was used (45 seconds at 2,000 RPM for 1 cycle) (Qiagen). The DNeasy PowerSoil extraction was performed using a QIAcube (Qiagen).

The hypervariable V4 region of the 16S rRNA gene was amplified to characterize the microbiome [40]. The universal 16S primers 515F/806R were used to PCR amplify the V4 region [41]. PCR was performed in triplicate, each reaction with a total volume of 25 µL. Each reaction contained 12.5 µL Phusion High-Fidelity PCR Master Mix with HF Buffer 2X concentration and 1 µL bovine serum albumin 20 mg/ml (New England BioLabs), 0.75 µL forward primer 10 µM, 0.75 µL reverse primer 10 µM, and 10 µL of DNA/molecular grade water. A total of 10 ng DNA was added per reaction. Thermocycler parameters were: denaturing step at 95 °C for 3 min, followed by 30 cycles of 95 °C for 45 s, 50 °C for 60 s, 72 °C for 90 s, and an extension step of 72 °C for 10 min. Subsequently, QIAxcel capillary electrophoresis (Qiagen) was utilized to assess presence of PCR product and determine the V4 band concentration for library pooling. PCR samples with similar concentrations (< 5 ng/µL, 5-10 ng/µL, > 10 ng/µL) were pooled together. Libraries clean-up was performed using GeneRead Size Selection kit (Qiagen). Libraries were sequenced on an Illumina MiSeq at the UConn Microbial Analysis, Resources, and Services facility. The reads were demultiplexed using the Illumina BaseSpace sequence hub and FASTQ files were downloaded for further data analysis.

Data was analyzed in R [42] using the dada2 package [43] to process data and generate Amplified Sequence Variants (ASV) and taxonomy tables. The forward and reverse reads were trimmed to 240 and 200 bp, respectively, and truncated using Q=11 and no Ns were allowed. The taxonomy was assigned to each ASV using silva_nr_v128. The phangorn package [44] was used to generate phylogenetic trees from ASV tables. Further analyses and sample filtering were performed in phyloseq [44]. Using the rarefy_even_depth function in phyloseq, microbiome count data were rarefied to sequencing depths of 1,000, 5,000, and 10,000 reads. Data were rarefied to these three levels to optimize microbiome sampling completeness, while trying to maximize sample sizes for analyses of effects related to host sex, host species, host guild, and geographical location. A sequencing depth of 1,000 reads was selected as minimum depth to retain the greatest number of samples for analyses, but this level discards a large amount of data from many samples and may include samples that are relatively poorly characterized. Increasing sequencing depth reduces the number of samples that meet the minimum requirements, resulting in reduced statistical power, but increases the relative completeness and number of rare ASVs included in samples. This represents a trade-off of statistical power for confidence in the characterization of the microbiome samples.

### Quantitative Analysis

Separately for oral and rectal samples from each host individual, we quantified microbiome biodiversity using four metrics based on ASVs: richness, Shannon diversity [46], Camargo’s evenness [47], and Berger-Parker dominance [48]. Hereafter, we refer to these metrics simply as “richness”, “diversity”, “evenness”, and “dominance”, and use “biodiversity” to refer to the general concept that comprises all 4 metrics. Each metric was expressed as Hill numbers, which are transformations based on relative abundances [48, 49]. Within the context of ASVs, Hill numbers are based on the relative number of reads that represent each ASV. Importantly, Hill numbers for all metrics are on the same scale (i.e. from 1 to richness) and in the same units (effective number of ASVs), which is defined as the number of equally abundant ASVs required to achieve the empirical value of a metric. Greater values for any Hill number represent greater biodiversity, including for dominance (i.e. larger values for Hill-transformed dominance indicate low dominance and greater biodiversity).

We used a 2-way analysis of variance (ANOVA) with type II sums of squares to evaluate effects of site (i.e. host population) and host sex for each host species that was represented by more than 1 population. Site and host sex were model I (fixed) treatment factors. *Artibeus jamaicensis* was captured at all three caves; *B*. *cavernarum*, *E*. *sezekorni* and *P*. *quadridens* were captured at Mata de Plátano and Río Encantado; and *M*. *redmani* was captured at Mata de Plátano and Aguas Buenas. For each host species without sufficient samples from multiple caves, but with samples for each sex, we used a general linear model (GLMM) to evaluate differences in microbiome biodiversity between males and females with host sex as a fixed effect and site as a random factor (i.e. model II treatment factor). Use of site as a random factor controlled for geographic variation to more powerfully evaluate differences between sexes in microbiome biodiversity

We used GLMMs to evaluate differences in microbiome biodiversity among host species for each guild (i.e. only among carnivorous species and only among herbivorous species) and between host guilds. Host species or host guild was a fixed effect and site was modeled as a random factor. Use of site as a random factor controlled for geographic variation to more powerfully evaluate species- or guild-level differences in microbiome biodiversity. For each GLMM that identified a significant difference in microbiome biodiversity between host species with a guild, we conducted a posteriori tests (Tukey’s test with a Holm-Šidák adjustment) to identify consistent differences between all possible pairs of host species. Because such a posteriori tests are less powerful than their associated GLMM and are protected in the sense that a posteriori tests were only executed when GLMMs were significant (α ≤ 0.05), we considered *P* ≤ 0.10 as evidence for significant pairwise differences.

For all analytical approaches, oral and rectal microbiomes were evaluated separately for each sequencing depth (i.e. 1,000, 5,000, and 10,000 reads) and analyses were conducted separately for each metric of biodiversity. For analyses based on host foraging guild, all host species were included to best represent variation associated with all carnivorous or herbivorous hosts. Because sample sizes decreased with increasing sequencing depth, the number of host species sometimes declined with greater sequencing depth.

### Accession Number

Sequencing data of V4 region of the 16S rRNA gene has been deposited in the NCBI Short Read Archive database under BioProject PRJNA602518 and accession numbers SRX7587313-7587772.

## RESULTS

Oral and rectal samples were collected from 331 individual bats, representing 10 species: 3 insectivorous mormoopids (*M*. *blainvillii*, *P*. *quadridens*, *P*. *parnellii*), 1 insectivorous vespertilionid (*E*. *fuscus*), 1 piscivorous noctilionid (*N*. *leporinus*), 2 frugivorous phyllostomids (*A*. *jamaicensis*, *S*. *rufum*), 1 nectarivorous phyllostomid (*M*. *redmani*), and 2 generalist herbivore phyllostomid (*B*. *cavernarum*, *E*. *sezekorni*). Samples were obtained from 10 bat species at Mata de Plátano (155 individuals), 9 species (all but *S*. *rufum*) at Río Encantado (101 individuals), and 6 species (75 individuals) at Aguas Buenas (*P*. *parnellii*, *N*. *leporinus*, *A*. *jamaicensis*, *M*. *redmani*, *B*. *cavernarum*, and *E*. *sezekorni*). As the bats were released after sampling, we used swabs to sample microbiomes, especially for the smaller species whose size made it challenging to extract sufficient amounts of microbial DNA for analysis. We obtained reasonable representation of the microbiomes (i.e. sequence depths > 1,000 reads) from less than half of those samples. Specifically, 136, 111, and 94 oral samples yielded sequencing depths of at least 1,000, 5,000, and 10,000 reads, respectively; and 157, 122, and 106 rectal samples yielded sequencing depths of at least 1,000, 5,000, and 10,000 reads, respectively.

Oral microbiomes comprised 2,114, 2,282, and 1,973 ASVs in samples with sequencing depths of 1,000, 5,000, and 10,000 reads, respectively. Rectal microbiomes comprised 2,986, 4,035, and 4,026 ASVs in samples with sequencing depths of 1,000, 5,000, and 10,000 reads, respectively. The reduction in number of ASVs between sequencing depths of 5,000 and 10,000 is due to the smaller number of samples available for analysis.

Bacteria represented over 98.8% of the ASVs in oral and rectal microbiomes from each host species. Archaea comprised the remainder of the microbiomes, occurring in the oral microbiomes of 8 of 10 host species (all but *N*. *leporinus* and *S*. *rufum*) and in the rectal microbiomes of 9 of 10 host species (all but *S*. *rufum*).

In aggregate, 37 and 36 phyla were identified from oral and rectal microbiomes, respectively; however, most of these taxa were represented by few ASVs and few reads of those ASVs. Only 16 and 14 phyla were represented by at least 5 ASVs from oral and rectal samples, respectively. Oral microbiomes were dominated by Actinobacteria (30.6% of all reads), Bacteroidetes (15.4%), and Firmicutes (29.2%). Actinobacteria was the most abundant phylum in oral microbiomes of 5 host species, including all 3 mormoopids, and 2 phyllostomids (a nectarivore and frugivore), whereas Firmicutes was the most abundant phylum in oral microbiomes of the remaining 5 host species, including the noctilionid, vespertilionid, and 3 phyllostomids (a frugivore and 2 generalist herbivores).

Rectal microbiomes were dominated by Actinobacteria (15.9% of all reads), Bacteroidetes (9.8%), Firmicutes (19.2%), and Proteobacteria (43.3%). The dominant phylum in rectal microbiomes (Proteobacteria) represented only 0.4% of oral microbiomes, but was the most abundant phylum in the rectal microbiomes of 9 host species, except for *E*. *fuscus*, for which Actinobacteria was the most abundant taxon.

Biodiversity was highly variable among individuals within each host species regardless of sequencing depth. Using sequencing depth of 1,000 as an example, maximum richness from an individual host for oral microbiomes was 3 to 38 (mean of 11) times greater than the minimum richness within host species. Similarly, maximum richness of rectal microbiomes from an individual host was 3 to 33 (mean of 9) times greater than the minimum within host species. Similar variation was observed within each host species for oral diversity (maximum 3 to 56 times that of the minimum, with a mean of 17) rectal diversity (maximum 3 to 57 times that of the minimum, with a mean of 20), oral evenness (maximum 2 to 55 times that of the minimum, with a mean of 17), rectal evenness (maximum 3 to 58 times that of the minimum, with a mean of 21), oral dominance (maximum 2 to 18 times that of the minimum, with a mean of 7), and rectal dominance (maximum 2 to 11 times that of the minimum, with a mean of 6).

The oral microbiome exhibited greater biodiversity than did the rectal microbiome in four host species, including 2 insectivorous mormoopids (*M*. *blainvillii* and *P*. *parnellii*) that harbor high microbiome biodiversity and 2 frugivorous phyllostomids (*A*. *jamaicensis* and E. *sezekorni*) that harbor low microbiome biodiversity (Table 1). In general, biodiversity of the more biodiverse microbiome (oral or rectal) was less than twice as great as its companion microbiome; however, *E*. *fuscus* (an insectivore) harbored rectal microbiomes that were more than 4 times as biodiverse as its oral microbiomes.

**Table 1.**
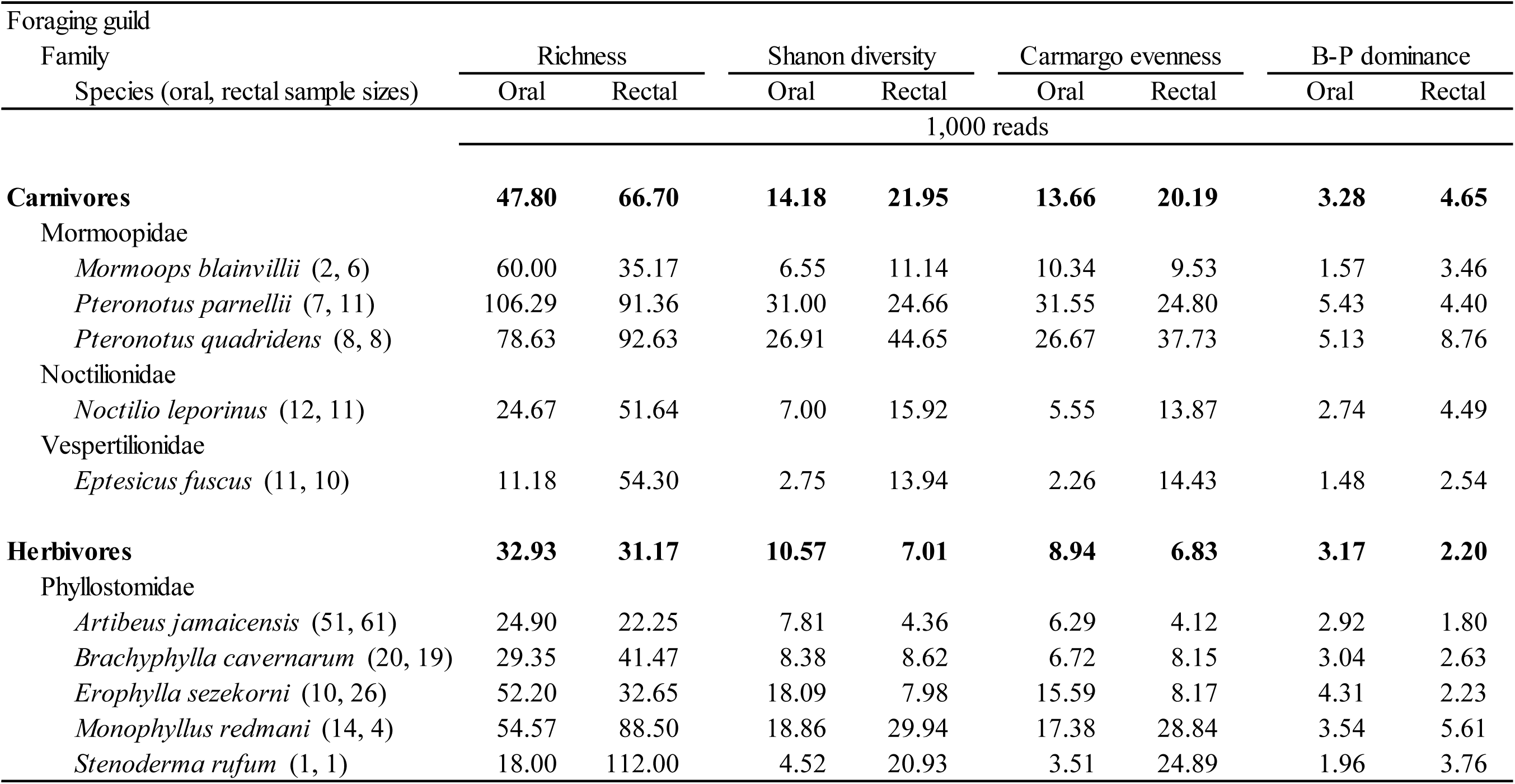
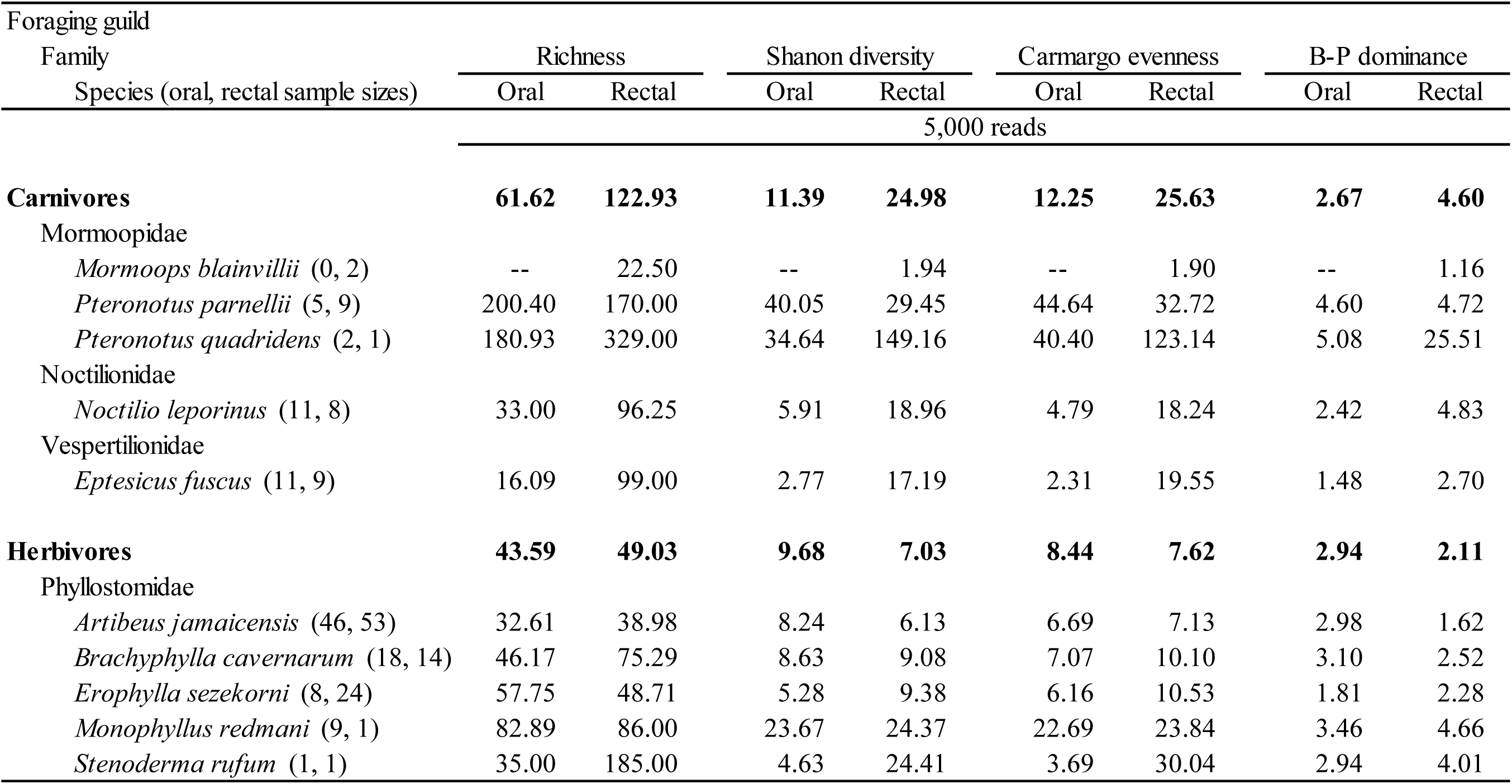
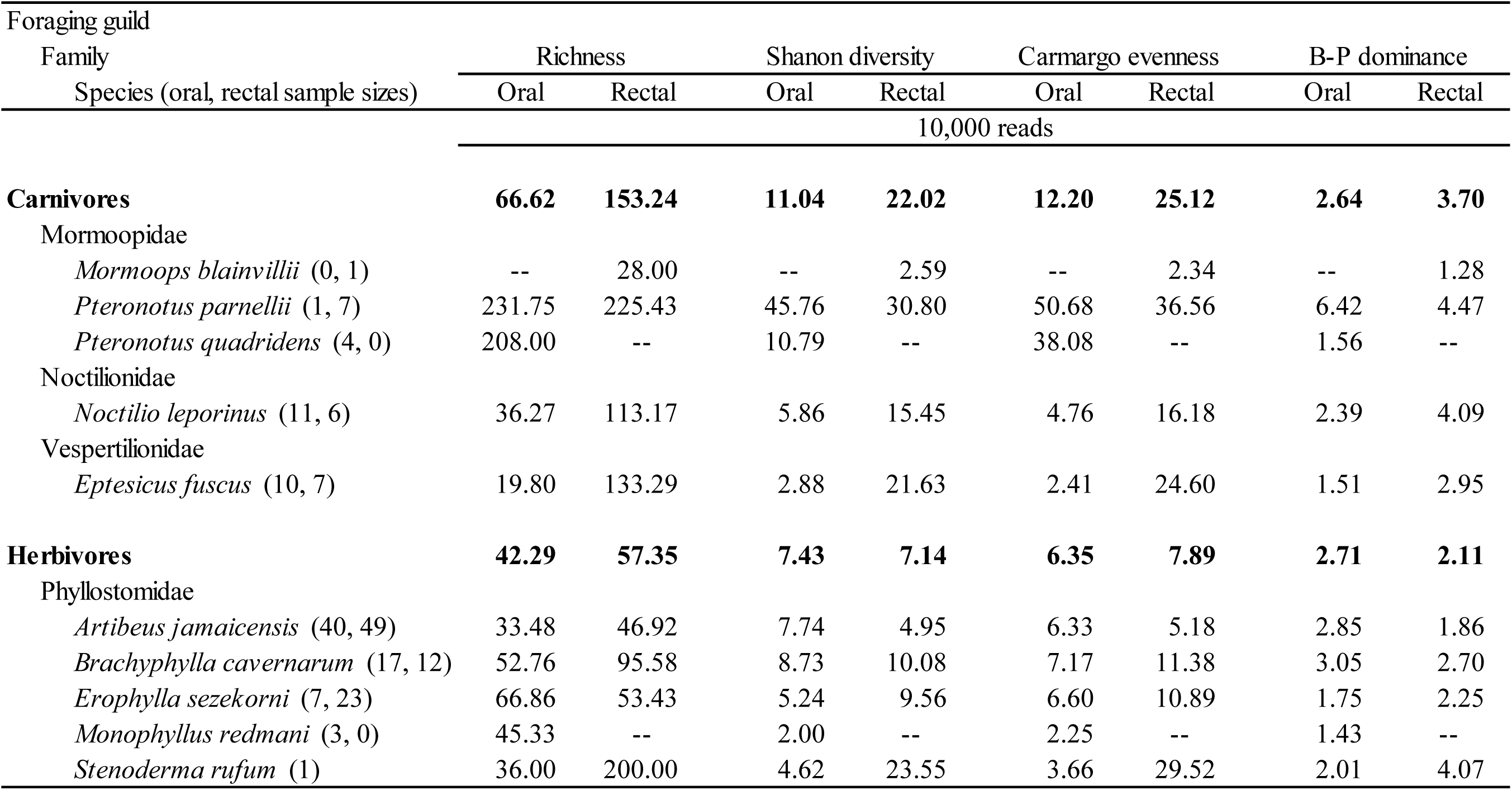
Mean biodiversity of oral and rectal microbiomes for each of 10 bat species in Puerto Rico as well as for all bats in each of two foraging guilds (carnivores and herbivores) regardless of species. Biodiversity was quantified using each of four metrics based on Amplified Sequence Variants (richness, Shannon diversity, Camargo evennes, Berger-Parker dominance) and expressed as Hill numbers. Guild-level values are bold

As expected, microbiome biodiversity increased as sequencing depth increased (Table 1). Insectivores had both the least (*E*. *fuscus*) and greatest (*Pteronotus* spp.) oral microbiome biodiversity. In contrast, frugivores (*A*. *jamaicensis*, *E*. *sezekorni*) had the least rectal microbiome biodiversity, and nectarivores (*M*. *redmani*) and insectivores (*Pteronotus* spp.) had the greatest rectal microbiome biodiversity (Table 1).

Host sex did not exhibit effects on oral or rectal microbiome richness (Table 2); however, at least one effect of sex on oral microbiome diversity, evenness, or dominance was found in *B*. *cavernarum* and on rectal microbiome diversity, evenness, or dominance in *M*. *blainvillii*, *P*. *quadridens*, and *A*. *jamaicensis* (Table 2). Consistent effects of site on oral microbiomes only manifested for *A*. *jamaicensis*, which had the greatest number of samples. At least one metric of rectal microbiome biodiversity responded to site for *A*. *jamaicensis* (richness), *E*. *sezekorni* (richness, evenness, and dominance), and *P*. *quadridens* (richness, diversity, evenness, and dominance) (Table 2). Oral and rectal microbiome biodiversity was greater from *A*. *jamaicensis* at Río Encantado than from *A*. *jamaicensis* at Mata de Plátano or Aguas Buenas (Fig. 3). For host species (*B*. *cavernarum*, *E*. *sezekorni* and *P*. *quadridens*) with sufficient sample sizes only at Río Encantado and Mata de Plátano, differences in oral microbiome biodiversity did not manifest between sites; however, rectal microbiome biodiversity was typically greater at Río Encantado than at Mata de Plátano (Fig. 4). Microbiome biodiversity from *M*. *redmani* did not differ between sites.

**Fig. 3.**
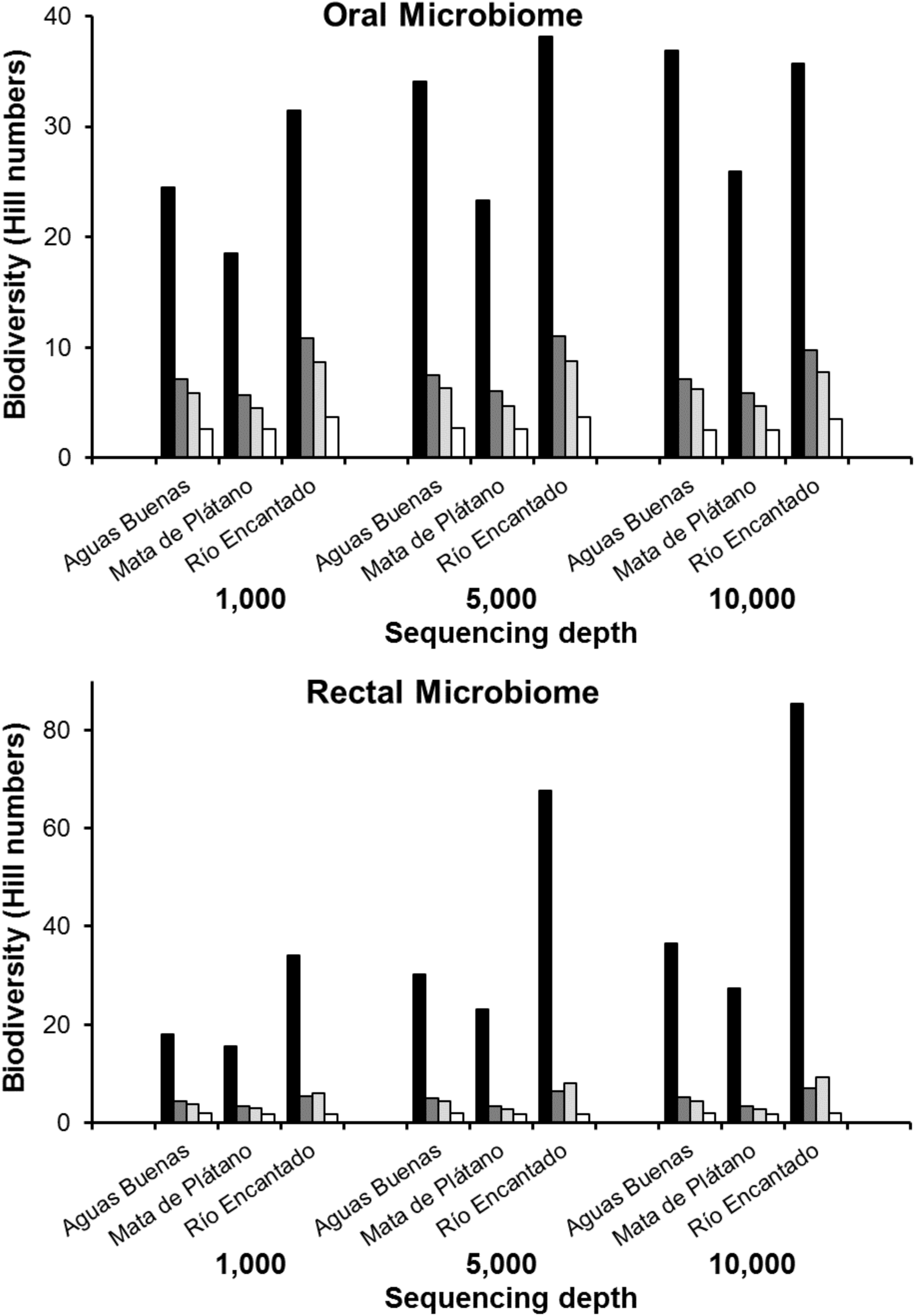
Aspects of biodiversity (richness, black bars; Shannon diversity, dark gray bars; Camargo’s evenness, light gray bars; Berger-Parker dominance, white bars) expressed as Hill numbers based separately on oral and rectal microbiomes for *Artibeus jamaicensis* at each of three sites (i.e. Aguas Buenas, Mata de Plátano, and Río Encantado) at sequencing depths of 1,000, 5,000, or 10,000 reads. In general, metrics of biodiversity for oral microbiomes were least at Mata de Plátano compared to other sites. For rectal microbiomes, only richness differed among sites, with Río Encantado exhibiting the greatest biodiversity. See Table 1 for details.

**Fig. 4.**
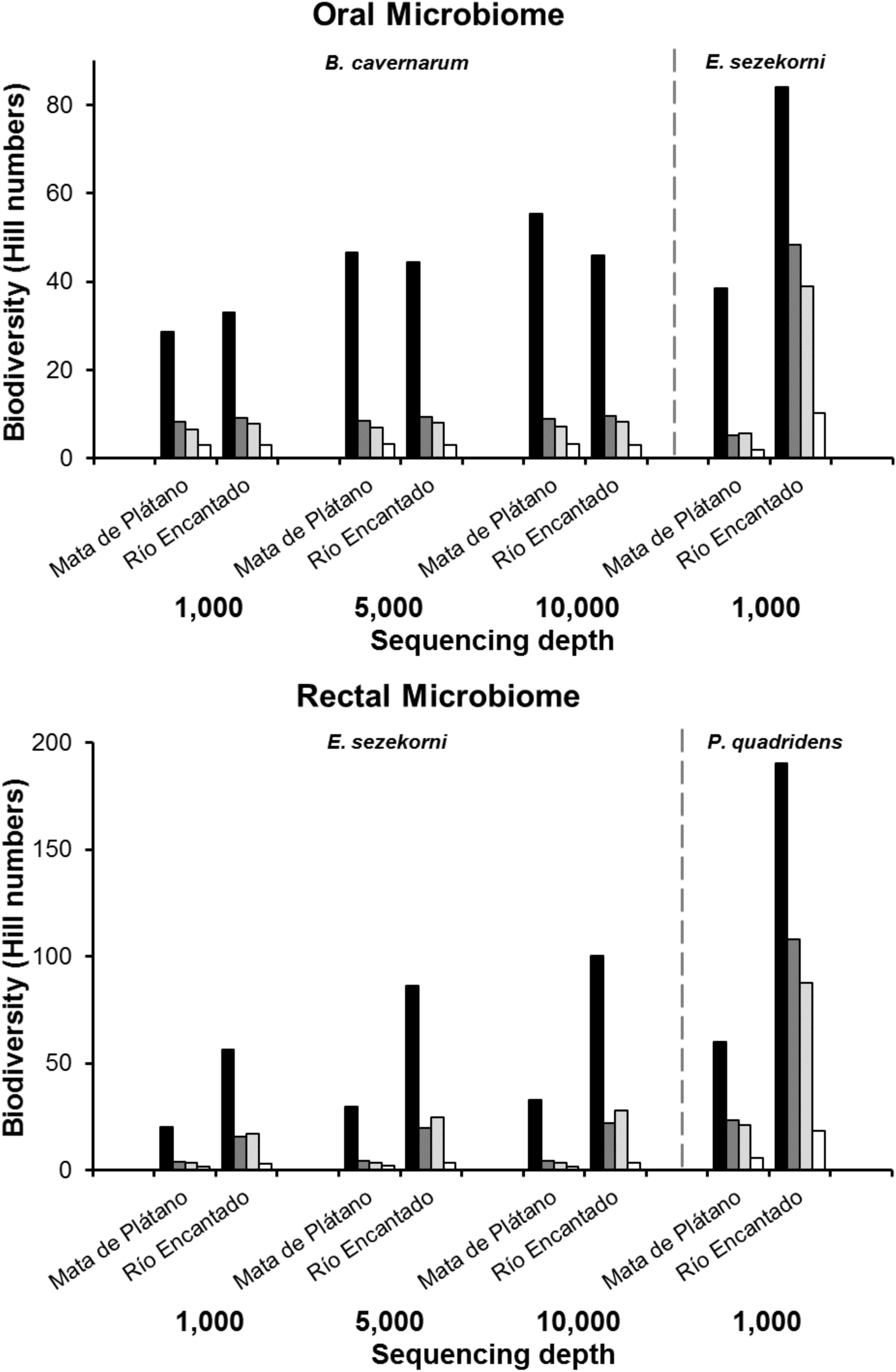
Aspects of biodiversity (richness, black bars; Shannon diversity, dark gray bars; Camargo’s evenness, light gray bars; Berger-Parker dominance, white bars) expressed as Hill numbers based separately on oral and rectal microbiomes for *Brachyphylla cavernarum*, *Erophylla sezekorni*, and *Pteronotus quadridens* from each of two sites (i.e. Mata de Plátano and Río Encantado) at sequencing depths of 1,000, 5,000, or 10,000 reads. No significant differences between sites characterized the aspects of biodiversity of the oral microbiome, whereas aspects of rectal microbiome biodiversity were generally greater at Río Encantado than at Mata de Plátano. See Table 1 for details.

**Table 2.**
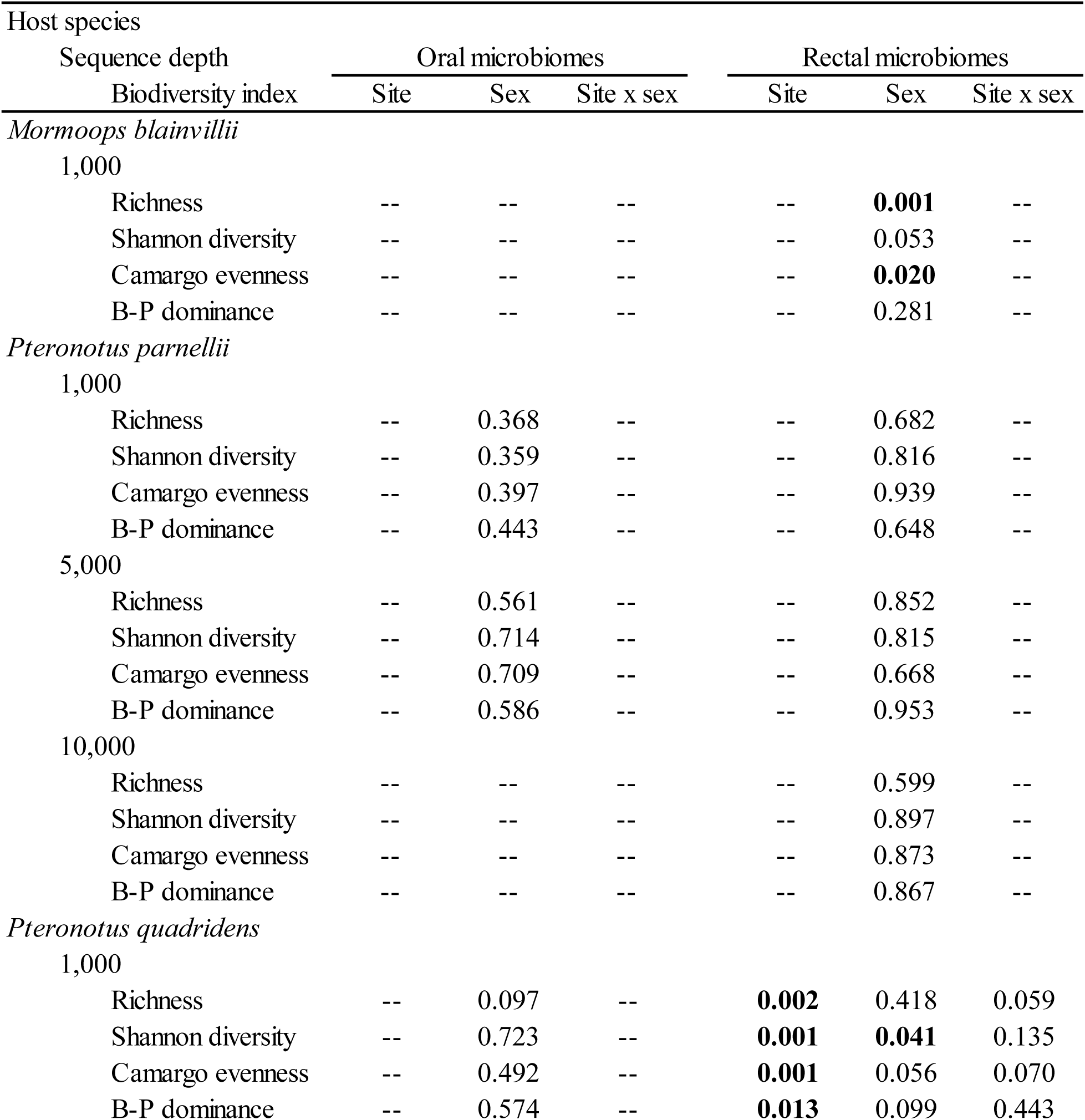
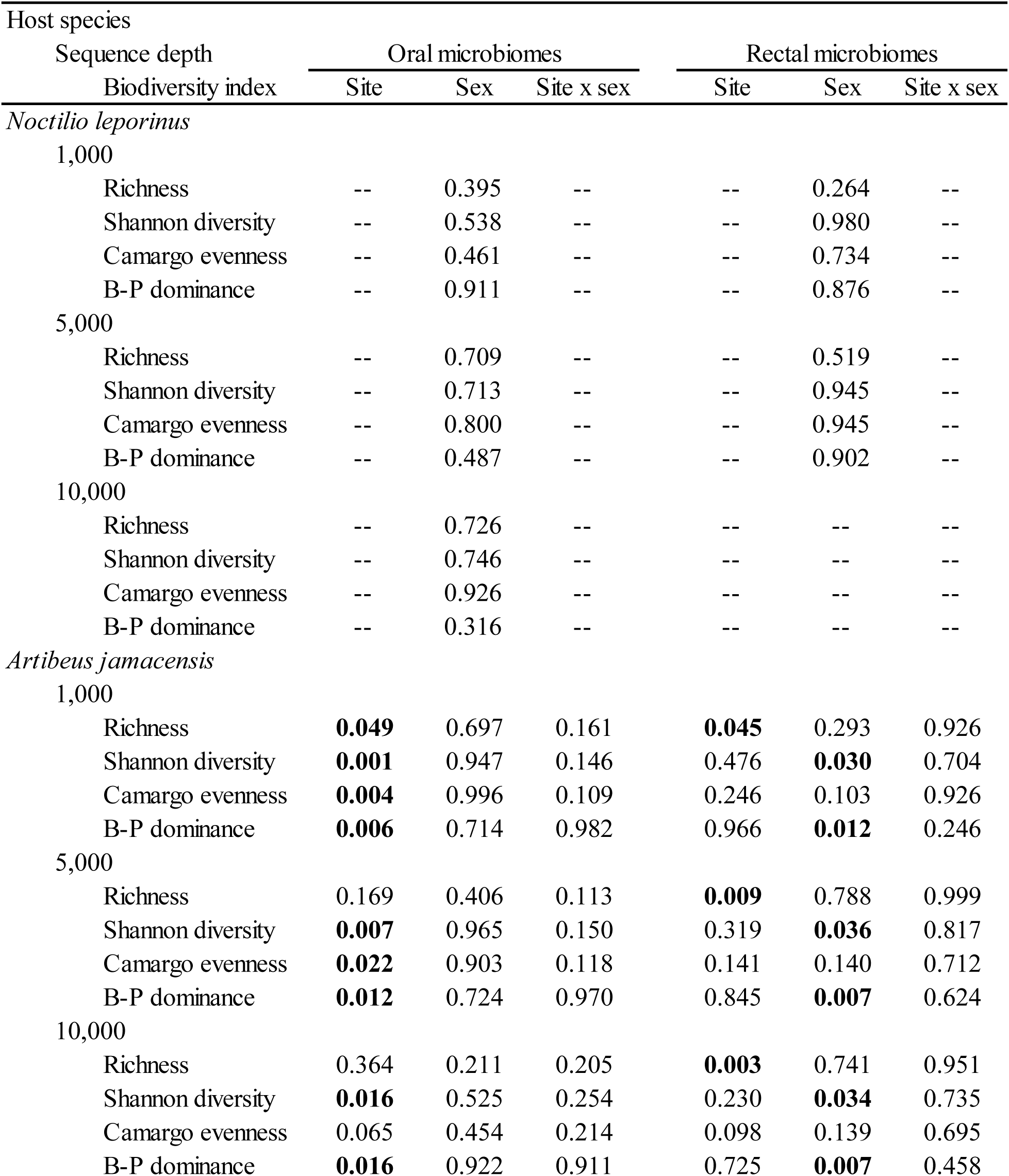
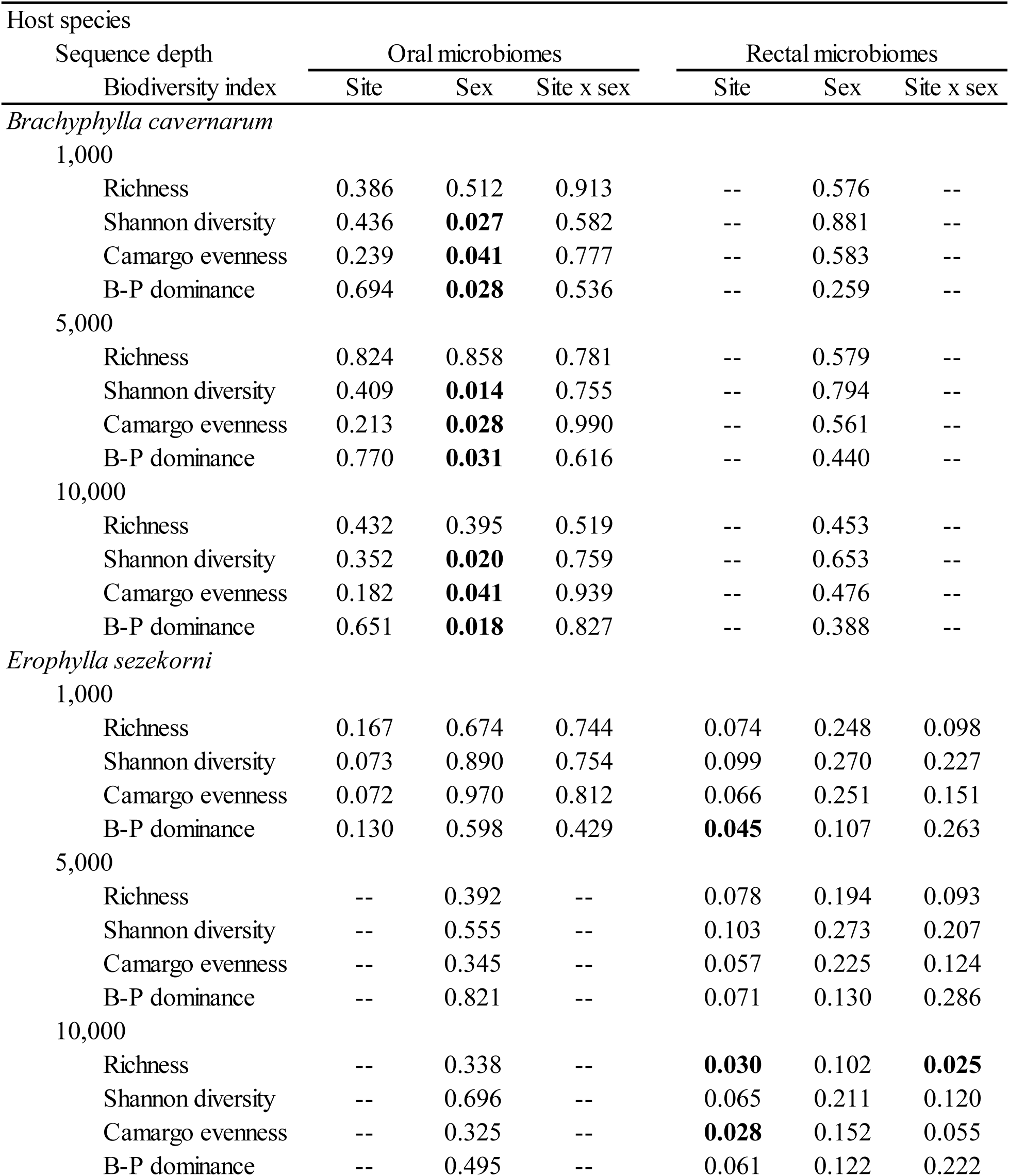
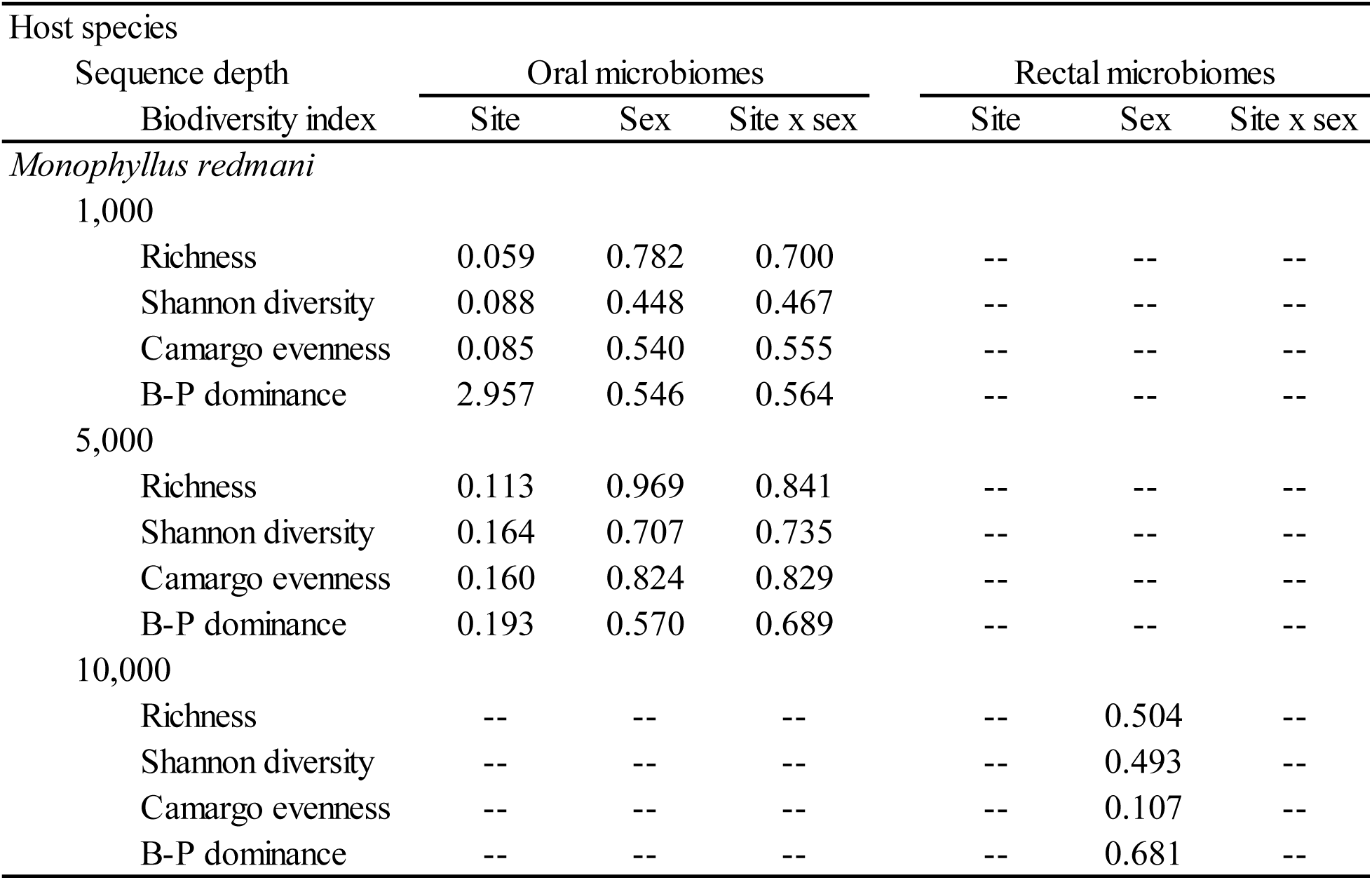
Results (*P* -values) of 1-way generalized linear mixed-effects models (for analyses of host sex only with site as a model II treatment factor) or 2-way analyses of variance with type II sums of squares (for analyses of site and host sex) evaluating the effects of site or host sex on microbiome biodiversity. Analyses were conducted separately for each combination biodiversity metric, sample type (oral or rectal), and sequencing depth. Significant results (*P* ≤ 0.05) are bold

Within each host guild, host species differed in oral microbiome biodiversity at each sequence depth; however, interspecific host differences in rectal microbiome biodiversity decreased with increasing sequence depth. We have stronger evidence for consistent species-specific differences in oral microbiome biodiversity within each guild than for species-specific difference in rectal microbiome biodiversity within each guild (Table S1). In contrast, no evidence suggests that guild-specific differences in oral microbiome biodiversity exist, whereas rectal microbiome biodiversity differed significantly between guilds (Table 3). Rectal microbiome biodiversity in carnivores was about twice that found in herbivores (Table 1; Fig. 5).

**Fig. 5.**
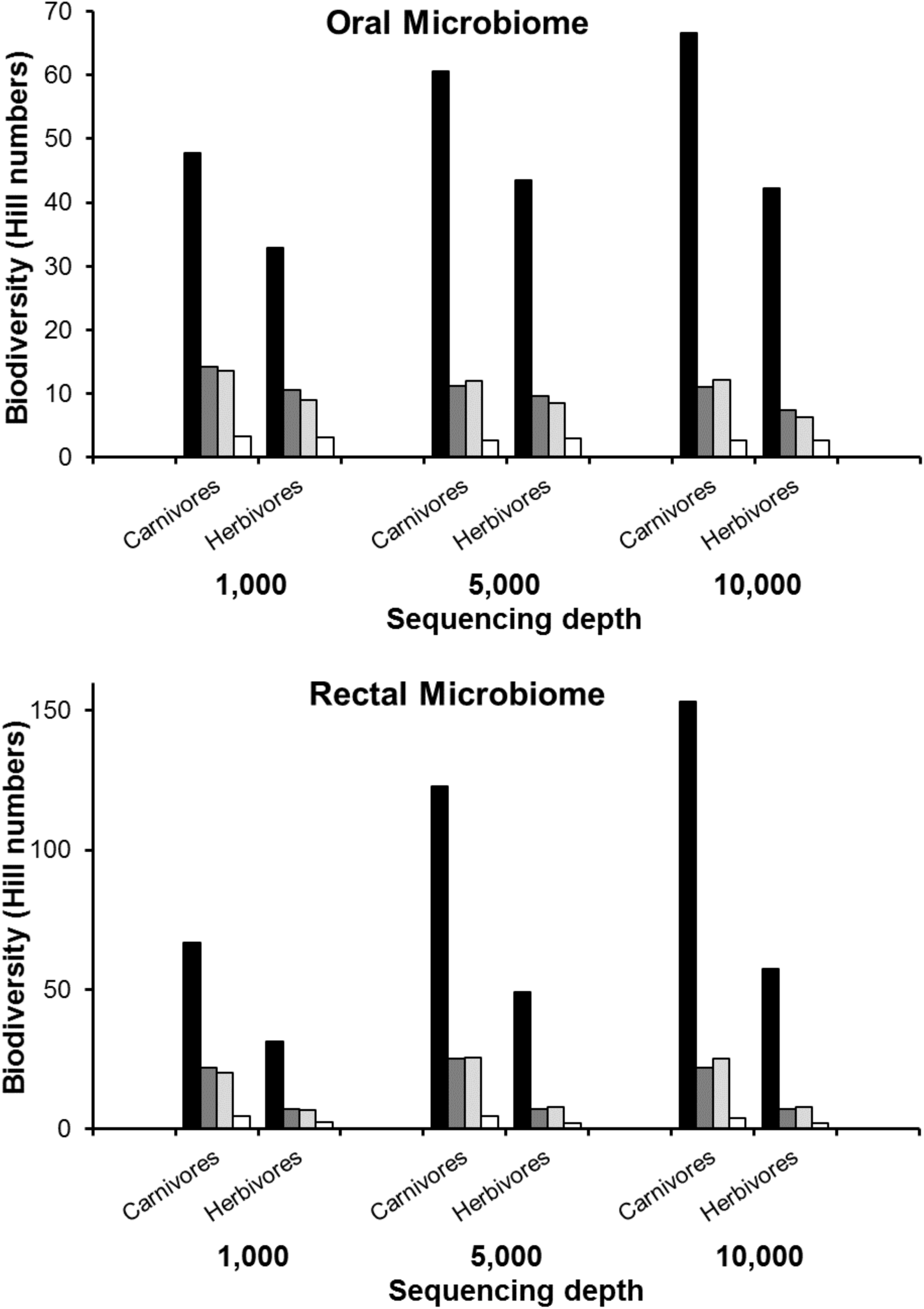
Aspects of biodiversity (richness, black bars; Shannon diversity, dark gray bars; Camargo’s evenness, light gray bars; Berger-Parker dominance, white bars) expressed as Hill numbers based separately on oral and rectal microbiomes for carnivorous and herbivorous bats at sequencing depths of 1,000, 5,000, or 10,000 reads. In general, metrics of biodiversity did not differ between foraging guilds for the oral microbiome, whereas metrics of biodiversity were significantly greater in carnivores than in herbivores for the rectal microbiome. See Table 1 for details.

**Table 3.** Results (*P* -values) of general linear mixed-effects models evaluating the effect of host species or host guild on microbiome biodiversity. Effect of host species was evaluated separately for each guild. Species and guild were model 1 treatment factors (i.e. fixed effects) and cave was a model II treatment factor (i.e. random effects). Analyses were conducted separately for each combination of biodiversity metric, sample type (oral or rectal), and sequencing depth. Significant results (*P* ≤ 0.05) are bold

| Sequencing depth | Comparison of species within guilds |  |  |  | Comparison between guilds |  |
| --- | --- | --- | --- | --- | --- | --- |
|  | Oral microbiome |  | Rectal microbiome |  | Oral microbiome | Rectal microbiome |
| Biodiversity index | Carnivores | Herbivores | Carnivores | Herbivores |  |  |
| 1,000 |  |  |  |  |  |  |
| Species richness | < **0.001** | **0.005** | 0.116 | < **0.001** | 0.061 | < **0.001** |
| Shannon diversity | < **0.001** | **0.048** | **0.009** | < **0.001** | 0.296 | < **0.001** |
| Camargo evenness | < **0.001** | **0.016** | **0.029** | < **0.001** | 0.120 | < **0.001** |
| B-P dominance | **0.002** | 0.567 | **0.006** | < **0.001** | 0.832 | < **0.001** |
| 5,000 |  |  |  |  |  |  |
| Species richness | < **0.001** | **0.008** | 0.143 | **0.011** | 0.188 | < **0.001** |
| Shannon diversity | < **0.001** | **0.005** | < **0.001** | 0.064 | 0.691 | < **0.001** |
| Camargo evenness | < **0.001** | **0.003** | **0.009** | 0.073 | 0.343 | < **0.001** |
| B-P dominance | < **0.001** | 0.122 | < **0.001** | **0.024** | 0.367 | < **0.001** |
| 10,000 |  |  |  |  |  |  |
| Species richness | < **0.001** | **0.034** | 0.363 | **0.005** | 0.094 | < **0.001** |
| Shannon diversity | < **0.001** | **0.015** | 0.589 | 0.095 | 0.829 | < **0.001** |
| Camargo evenness | < **0.001** | 0.205 | 0.534 | 0.075 | 0.266 | < **0.001** |
| B-P dominance | < **0.001** | **0.025** | 0.575 | **0.049** | 0.093 | < **0.001** |

## DISCUSSION

Considerable intraspecific variation characterized microbiome biodiversity, even after controlling for geography or sex of the host individual. These results mirror those for fecal and gastrointestinal microbiomes from vespertilionid bats of Slovenia [51] and from emballonurid, molossids, mormoopid, phyllostomid, and vespertilionid bats from Costa Rica [14], for which variation among conspecific hosts was high. This suggests that studies relying on a few samples per host species [9, 13] do not accurately capture variation in microbiome biodiversity or composition that naturally occurs within populations. Consequently, ecological conclusions based on such small samples may not be reliable, as estimates of biodiversity may not be accurate (especially richness) and statistical power to detect differences in any metric would be quite low.

Greater microbiome biodiversity in a host species could arise in two ways: 1) an increase in the number of Phyla or Classes of microbes found in the microbiome, or 2) an increase in the number of ASVs that belong to the same Phyla or Classes of microbes (i.e. not an increase in higher level taxonomic biodiversity). For both oral and rectal microbiomes, the latter scenario occurred. Host species with greater microbiome biodiversity (e.g. *P*. *parnellii*, *P*. *quadridens*, *M*. *redmani*) typically harbored more ASVs belonging to the same Phyla as those present in hosts with low microbiomes biodiversity. Archaea richness and Bacteria richness at the host species level (i.e. data combined for all hosts belonging to the same species) were highly correlated (oral, *R* = 0.928, *P* < 0.001; rectal, *R* = 0.690; *P* = 0.027). Similarly, pairwise correlations between richness values of different Phyla at the host species level indicate positive associations predominate (i.e. an increase in microbiome richness is associated with an increase in richness for most of the Phyla present). In oral microbiomes, 70% of pairwise correlations of Phylum relative abundances at the host species level were strongly positive (R > 0.50), and in rectal microbiomes 56% of pairwise correlations of Phylum abundances at the host species level were strongly positive (R > 0.50). Such correlations also characterize vespertilionid, rhinolophid, and miniopterid bats from Slovenia [51].

### Effects of host sex

Host sex could affect microbiome biodiversity of bat hosts due to differences in social organization or diet. Harems, comprising several adult females with 1 adult male, are common social structures for noctilionid [52] and phyllostomid bats [53], whereas maternity colonies, comprising adult females and their offspring, are common in mormoopid [54] and vespertilionid [55] bats. In contrast, most adult males are solitary in both of these social systems. In addition, the diets of male and female bats differ during some seasons, especially during periods of pregnancy and lactation when females target food sources that are higher in energy and protein [56, 57]. Despite sampling during the reproductive season, when these sex-based ecological differences manifest most strongly, we found little evidence of differences between sexes based on oral or rectal microbiome biodiversity (Table 2). When evidence of differences in microbiome biodiversity did manifest (i.e. in oral microbiomes of *B*. *cavernarum* and rectal microbiomes of *A*. *jamaicensis*), those differences were in the relative abundances of the ASVs (diversity, evenness, or dominance) in the microbiomes and not in the number of ASVs (richness). Fecal microbiomes from 12 species of vespertilionid bat from Slovenia failed to reveal differences between the sexes [51].

### Effects of geographical location

Despite the potential for environmental factors (e.g. roost environment, abundance and diversity of hosts in the roost) to affect oral microbiome biodiversity, only *A*. *jamaicensis* exhibited site-specific differences in oral microbiome biodiversity (Table 2; Figs 2 & 3). These differences may be related to population size or to host species diversity in associated roosts. Oral microbiomes from *A*. *jamaicensis* in Río Encantado had the greatest biodiversity, whereas those from Mata de Plátano (Larva Cave) had the lowest biodiversity. The population of *A*. *jamaicensis* at Río Encantado was greater than at other locations, and especially compared to Mata de Plátano. Moreover, the number of bats and bat species was much greater at other caves than at Larva, where *A*. *jamaicensis* roosts at Mata de Plátano. Of course, populations sizes differed among sites for other host species without significant differences in oral microbiome biodiversity. This suggests that host abundance may not be the major factor determining oral microbiome biodiversity. In general, intraspecific variation in oral microbiome composition and biodiversity is high and may rival interspecific variation.

Rectal microbiomes of each host species exhibited site-specific variation in biodiversity (Table 2; Figs 2 & 3). In *A*. *jamaicensis*, rectal microbiomes exhibited patterns similar to those observed for oral microbiomes, with greater biodiversity associated with larger populations from roosts with greater bat species richness. In contrast, rectal microbiomes from *E*. *sezekorni* and *P*. *quadridens* exhibited greater biodiversity from Río Encantado than from Mata de Plátano, with the former harboring fewer individuals than the latter. Host abundance or biodiversity may not have direct effects on microbiome biodiversity, but may serve as proxies for important ecological factors. For example, bat abundance or diversity may be related to the diversity or abundance of dietary items or habitat types used by resident bats, and the diversity of diet or habitat may influence spatial patterns of microbiome biodiversity. Alternatively, microbiome biodiversity within sites may represent legacies or factors such as the effects of hurricane-induced disturbances on bat populations and communities [39]. Although confident identification of causal mechanisms that drive spatial variation in microbiome biodiversity is challenging and beyond the scope of this study, our results strongly suggest that spatial variation must be considered when evaluating aspects of microbiome biodiversity, especially for rectal microbiomes.

### Effects of host species or guild on biodiversity of oral microbiomes

Within each host guild, species-specific differences characterized biodiversity of oral microbiomes. In contrast, guild-specific differences did not characterize oral microbiomes (Table 3). This combination of results indicates that oral microbiome biodiversity is unrelated to host diet for Puerto Rican bats. For carnivores, nearly all pairwise comparisons of oral microbiome biodiversity between host species were significant (Table S1), suggesting distinct oral microbiome biodiversity for each carnivorous species. In contrast, pairwise differences in oral microbiome biodiversity among herbivorous bat species were primarily driven by differences between *M*. *redmani* (most diverse oral microbiome) and other herbivores.

Patterns of oral microbiome biodiversity may be structured by processes similar to those of microbiomes from other mucosal surfaces (e.g. nose, mouth, vagina, lungs, gastrointestinal tract). The microbiome of the mucosal lining of the intestines directly interacts with the host immune system through receptors in the intestinal epithelia [58]. The direct sampling of the intestinal mucosa showed a strong relationship between intestinal microbiome composition and host phylogeny in Belizean bats [14]. The species-specific biodiversity observed for oral microbiomes within each guild of bats in Puerto Rico likely represents a similar co-evolutionary association between hosts and their microbiomes. The carnivores represent 3 families of bats (Mormoopidae, Vespertilionidae, and Noctilionidae), which likely contribute to the preponderance of significant pairwise differences in the biodiversity of oral microbiomes. In contrast, the lower frequency of pairwise differences in oral microbiome biodiversity among herbivorous species likely arises because they represent a single family (Phyllostomidae) of bats.

### Effects of host species or guild on biodiversity of rectal microbiomes

Species-specific differences with host guilds exhibited 2 patterns: (1) species-specific differences were more consistent at lower sequencing depths than at greater sequencing depths and (2) species-specific differences were observed more consistently between species of herbivore than between species of carnivore (Table 3). In contrast, consistent differences in biodiversity occurred between the rectal microbiomes of carnivorous and herbivorous foraging guilds (Table 3). In concert, these results suggest that the biodiversity of rectal microbiomes is related to host diet. Regardless of metric, the biodiversity of rectal microbiomes of carnivorous bats (mostly insectivores) were nearly twice as great as those from herbivorous (mostly frugivores) bats (Table 1; Fig. 5). Importantly, the lack of species-specific differences in biodiversity within host foraging guilds in some cases does not suggest that the composition of rectal microbiomes does not differ among species within a guild. Indeed, microbe composition may differ among host species within a guild, with different microbe taxa performing the same function in different host species. However, the number of microbial taxa that a host supports may be contingent on the general diet of the host species (i.e. the number and kinds of functions a host requires of its microbiome). This is consistent with findings from a soil and plant microbiome assembly experiment in which metacommunities contained fixed fractions of coexisting families that were determined by the available carbon source [59]. Despite consistent higher level (Familial) structure, these assembled microbiomes exhibited great variation in taxonomic composition with the same functions performed in each microbiome but done so by different confamilial taxa.

Microbiomes associated with the digestive system from insectivorous bats are more biodiverse than their herbivorous counterparts in Guatemala [9], Mexico [13], and Puerto Rico (Tables 1 and 3). Greater microbiome biodiversity in carnivorous bats contrasts with theory based on the study of a wide array of mammals (e.g. ruminants, primates, carnivores). Three general predictions have been postulated [60]: (1) herbivorous hosts should have the most complex gut morphologies and most diverse microbiomes; (2) carnivorous hosts should have the most simple gut morphologies and the least biodiverse microbiomes; and (3) omnivorous hosts should have intermediate levels of gut complexity and microbiome biodiversity. Regardless of diet, all bats have shorter intestines and shorter food-retention times compared to similarly sized non-volant mammals as an adaptation for flight [32, 61]. Nonetheless, herbivorous bats still have slightly larger intestines than do carnivorous counterparts of similar size [62]. In contrast to non-volant herbivorous mammals that feed primarily on leaves or grass, herbivorous bats generally consume nectar and fruits that are poor sources of energy and nutrients, and that primarily contain simple sugars and carbohydrates, resulting in brief retention times (i.e. < 60 minutes) [63, 64]. Moreover, herbivorous bats rely on paracellular absorption for > 70% of their glucose absorption, which may explain why these bats have relatively depauperate rectal microbiomes [32, 65]. In contrast, the high protein, lipid, and nutrient content of insectivorous diets may result in high microbiome biodiversity due to the variety of carbon and energy sources available [13].

## Conclusions

High variation in microbiome diversity among individuals of the same species suggests that individual-level host traits may affect the associated microbiome. Although initial, descriptive studies may provide new insights from few samples per host species, research designed to explore the ecological dynamics of microbiomes should account for such variation by increasing the number of samples collected from host populations. Despite effects of host ecology and evolutionary history on microbiome biodiversity, microbiome composition and biodiversity are also affected by spatial phenomena, primarily via host-environment interactions. Future work should investigate the roles of environmental factors that mediate microbiome biodiversity to decouple these effects from those associated with host ecology and evolution.

## Acknowledgments

This work was supported by a Microbiome Research Seed Grant to MRW and JG from the Office of the Vice President of Research at the University of Connecticut. In addition, SJP and MRW were supported by the National Science Foundation (DEB-1546686 and DEB-1831952) and by the Center for Environmental Sciences and Engineering at the University of Connecticut. We gratefully acknowledge our team of field volunteers, Armando Rodriguez-Duran for logistical help, and Para la Naturaleza for providing access to the Rio Encantado field site.

